# Genome sequence of the aurodox-producing bacterium *Streptomyces goldiniensis* ATCC 21386

**DOI:** 10.1101/2021.06.01.446637

**Authors:** Rebecca E. McHugh, John T. Munnoch, Andrew J Roe, Paul A. Hoskisson

## Abstract

We report the genome sequence of *Streptomyces goldiniensis* ATCC 21386, a strain which produces the anti-bacterial and anti-virulence polyketide, aurodox. The genome of *S. goldiniensis* ATCC 21386 was sequenced using a multiplatform hybrid approach, revealing a linear genome of ~10 Mbp with a G+C content of 71 %. The genome sequence revealed 36 putative biosynthetic gene clusters (BGCs), including a large region of 271 Kbp that was rich in biosynthetic capability. The genome sequence is deposited in DDBJ/EMBL/GenBank with the accession number PRJNA602141.

## Introduction

Isolated from soil collected in Bermuda in 1973, a novel strain of *Streptomyces* was found to produce the anti-Streptococcal natural product (X-5108 = aurodox) and was informally named *S. goldiniensis* var. *goldiniensis* [1]. Despite the strain being named *S. goldiniensis* var. *goldiniensis* by Berger et al., [1] there is no formal description of this strain in the literature and it does not appear in the List of Prokaryotic names with standing in nomenclature [2].

Originally identified its antibacterial activity, aurodox [1] has also found utility as a widely used growth promoting compound in poultry [3]. More recently it has attracted attention for its anti-virulence activity, blocking virulence by inhibition of the Type III Secretion System (T3SS) in Enterohaemorrhagic *Escherichia coli* (EHEC) [4]. Subsequent work this inhibitory effect is mediated through the down-regulation of the Type III Secretion System master regulator Ler [5].

Whilst *S. goldiniensis* var. *goldiniensis* is known for the production of aurodox, it is a common feature of *Streptomyces* genomes to encode many more secondary metabolites than can observed during laboratory culture [6]. The vast repository of natural product biosynthetic gene clusters (BGCs) contained within the genomes of *Streptomyces* means that genome mining, has become the mainstay of researchers looking to prioritize BGCs for further study [6, 7]. Moreover, the increasing amount of genome sequence data of natural product producing strains facilitates evolutionary studies of biosynthesis and can inform on synthetic biology strategies for developing novel molecules and improving production of current molecules [6–8]. Here we describe the multi-platform genome sequencing of *Streptomyces goldiniensis* ATCC 21386 which is deposited in DDBJ/EMBL/GenBank with the accession number PRJNA602141.

## Results & Discussion

### Genome features of *Streptomyces goldiniensis* ATCC 21386

The linear genome of *S. goldiniensis* ATCC 21386 was sequenced using a hybrid-approach of Illumina, PacBio and Oxford Nanopore to generate a high-quality draft genome (Genbank Bioproject PRJNA602141). Using a combined assembly approach with SPAdes [9]the data from all three platforms allowed the overall genome size to be estimated at 10,005,022 bp, in nine contigs, with an N50 of 9,950,726 bp. The draft genome of *S. goldiniensis* is predicted to have a total of 9925 protein coding genes, along with 81 tRNAs and five rRNA operons (**Figure 1**). Absence of genes encoding plasmid replication machinery (Par proteins) on the eight minor contigs suggest that they do not represent plasmids. Pulse-filed Gel electrophoresis of total DNA extractions from *S. goldiniensis* also indicates the absence of plasmids in this strain.

**Figure 1:**
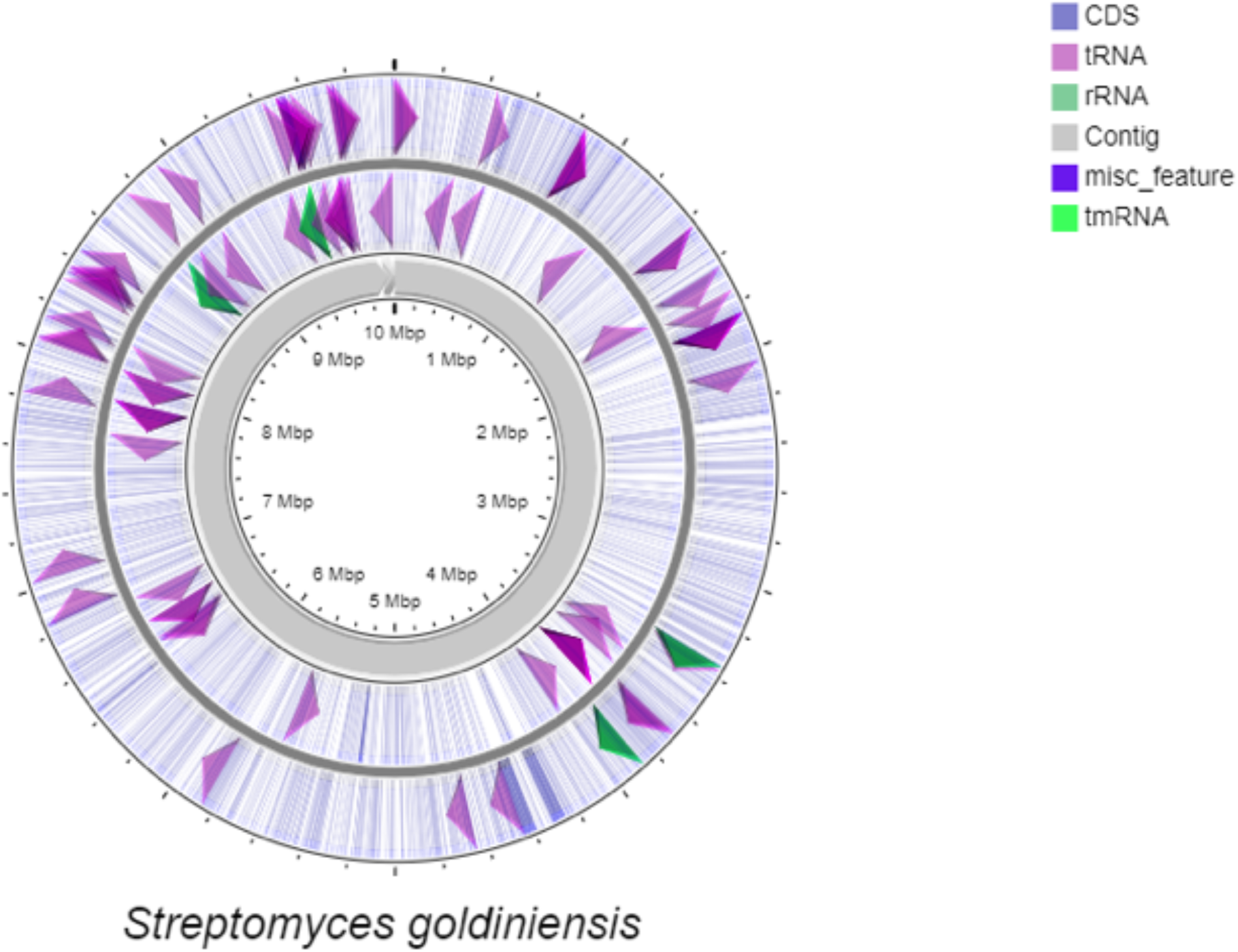
Circular representation of the linear *Streptomyces goldiniensis* genome generate by CGviewer [19]. Map was built from a .gbk file generated by Prokka. The outer most circle represents coding sequences on the major strand, the inner circle represents coding sequences on the minor strand. Grey arrows in the centre of the figure depict the lengths of the contigs in the final assembly.

### The *S. goldiniensis* genome is rich in natural product biosynthetic gene clusters

*Streptomyces* bacteria are renowned for their ability to synthesis a wide range of natural products, many of which have found utility in human medicine [7]. Despite many strains being identified through the production of a single metabolite, the genomes of *Streptomyces* often encode a number of additional biosynthetic gene clusters (BGCs) that are not expressed under laboratory conditions, the so called ‘silent BGCs’[6]. Using the antiSMASH pipeline [10] the genome of *S. goldiniensis* ATCC 21386 was mined. A total of 36 putative BGCs were identified (**Table 1**), including five putative polyketide synthase (PKS) containing BGCs, eight non-ribosomal peptide synthase (NRPS) BGCs and nine putative terpene BGCs. The genome encodes BGCs that are highly conserved in the genomes of *Streptomyces* species such as geosmin, desferrioxamine and melanin [7].

**Table 1:** Predicted Biosynthetic Gene Clusters of *S. goldiniensis* using antiSMASH [11].

| Region | Type | Region | Most similar | % similarity |
| --- | --- | --- | --- | --- |
| 1 | Type II PKS | 32,278 - 104,355 | Spore pigment | 83 |
| 3 | Lanthipeptide, lassopeptide | 486,431 - 514,192 | Citrulassin E (ripp) | 100 |
| 4 | Siderophore | 2,159,777 - 2,170,184 | NA | 0 |
| 5 | Terpene | 2,487,747 - 2,508,882 | Lipopeptide | 4 |
| 6 | NRPS, tfuA related | 2,543,021 - 2,606,769 | Diisonitrile antibiotic SF2768 | 66 |
| 7 | Bacteriocin | 2,705,875 - 2,716,826 | NA | 0 |
| 8 | Terpene | 2,777,350 - 2,799,499 | Geosmin | 100 |
| 9 | NRPS-like | 2,818,277 - 2,858,728 | S56P1 | 11 |
| 10 | NRPS-like, siderophore | 3,085,338 - 3,126,119 | Paulomycin | 13 |
| 11 | NRPS-like | 3,541,716 - 3,582,164 | Octacosamicin | 12 |
| 12 | Terpene | 3,625,827 - 3,643,980 | NA | 0 |
| 13 | Indole | 4,158,688 - 4,181,228 | Terfestatin | 28 |
| 14 | bacteriocin,bottromycin,NRPS,transAT-PKS, Type I PKS, hglE-KS,lassopeptide | 4,213,370 - 4,484,508 | Kirromycin/bottromycin | 40 |
| 15 | NRPS-like | 4,599,349 - 4,657,342 | Echosides | 17 |
| 16 | Terpene | 4,783,720 - 4,802,511 | NA | 0 |
| 17 | Linardin | 5,046,397 - 5,067,017 | Pentostatine | 17 |
| 18 | Siderophore | 5,085,773 - 5,100,346 | NA | 0 |
| 19 | NRPS, melanin | 5,167,798 - 5,232,264 | Scabichelin | 100 |
| 20 | Terpene | 5,257,095 - 5,277,644 | NA | 0 |
| 21 | NRPS, Type I PKS | 5,284,180 - 5,346,580 | Herboxidiene | 4 |
| 22 | Terpene | 5,480,336 - 5,505,711 | Isorenieratene | 100 |
| 23 | Type II PKS | 5,770,343 - 5,856,353 | Fluostatins | 60 |
| 24 | Lanthipeptide, bacteriocin | 6,153,412 - 6,179,099 | Informatipeptin | 100 |
| 25 | NRPS-like, Type I PKS | 6,410,852 - 6,457,973 | Marineosin | 45 |
| 26 | Terpene | 6,507,863 - 6,532,462 | Hopene | 100 |
| 27 | Ectoine | 6,852,064 - 6,861,735 | Ectoine | 100 |
| 28 | Ladderane | 6,937,157 - 6,978,500 | Colabomycin | 11 |
| 29 | Type III PKS, Terpene | 7,764,486 - 7,805,661 | Merochlorin | 12 |
| 30 | Melanin | 8,189,939 - 8,200,406 | Melanin | 80 |
| 31 | Siderophore | 8,344,177 - 8,355,949 | Desferrioxamine A | 83 |
| 32 | Ectoine | 8,471,171 - 8,481,599 | Ectoine | 75 |
| 33 | Terpene | 9,170,031 - 9,191,797 | NA | 0 |
| 34 | Bacteriocin | 9,838,044 - 9,848,331 | NA | 0 |
| 35 | Terpene | 9,968,421 - 10,005,017 | NA | 0 |

One region of the *S. goldiniensis* genome was found to be particularly rich in BGCs (position 4,213,370 - 4,484,508; 271 kb), encoding a putative Bottromycin A2-like molecule, an 87 kb region that possesses genes likely to encode a hybrid PKS/NRPS, highly similar to the kirromycin BGC [11] which has recently been shown to encode aurodox [12]. Immediately downstream of the aurodox BGC is a gene cluster with homology to glycolipid synthase-like PKS containing BGCs, a cluster which is 100 % identical to the macrolide concanamycin A and a putative lassopeptide-encoding gene cluster (**Table 1; Figure 2**). To confirm that this BGC-rich region around the aurodox BGC was indeed a supercluster, rather than an artefact of genome assembly, primer pairs were designed that spanned the junction between the cluster upstream (putative bottromycin A2) and downstream (glycolipid synthase-like PKS containing BGC) of the aurodox cluster. PCR and sequencing of the regions spanning the BGCs confirmed the organisation of the gene clusters around aurodox (**Figure 2**).

**Figure 2:**
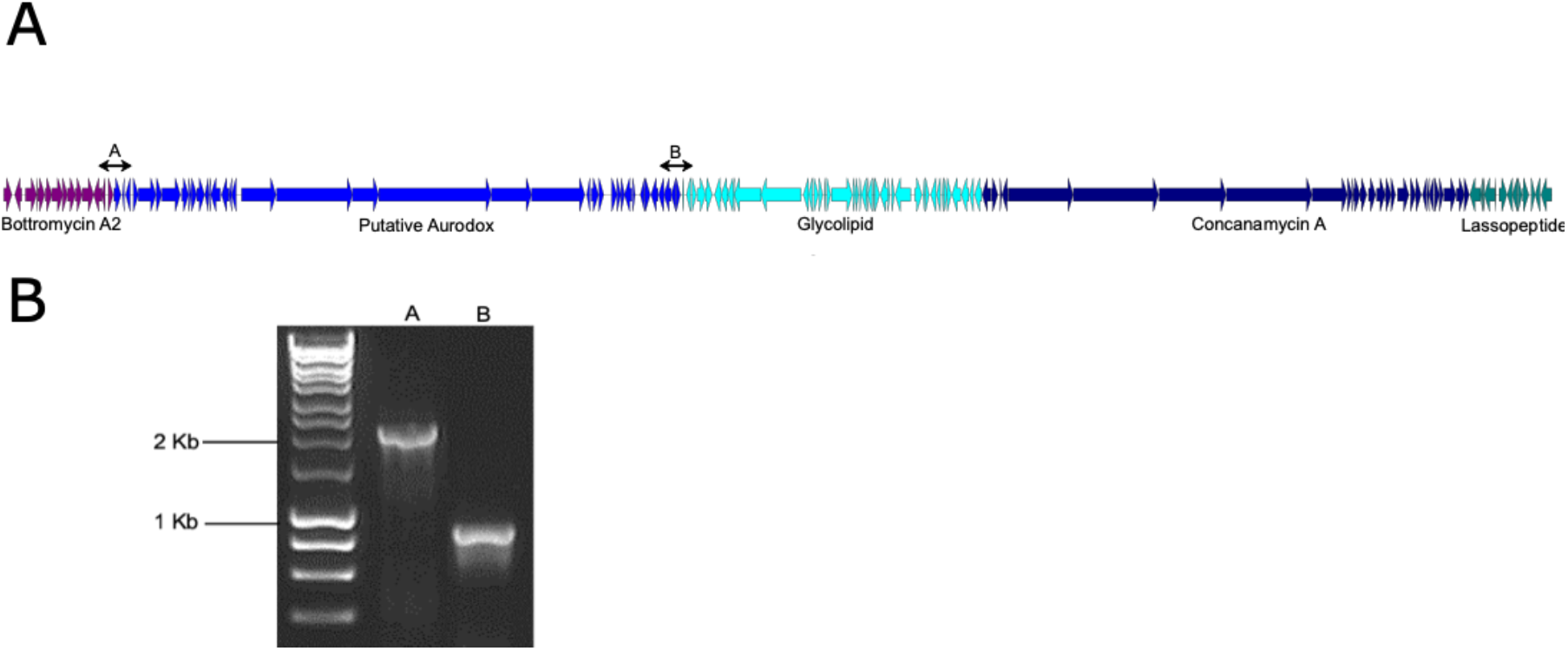
A: Representation of the 271 kbp putative aurodox supercluster. Gene cluster boundaries were predicted by AntiSMASH and schematic diagram was generated by clinker. From left to right, arrows coloured purple represent Bottromycin A2-associated genes, blue arrows represent the putative aurodox BGC, cyan arrows represent predicted glycolipid-synthase genes, navy blue arrows represent concanamycin A-associated genes and green arrows indicate putative genes involved in lassopeptide biosynthesis. **B:** Primer pair binding sites are indicated by ‘A’ and ‘B’ on the diagram and their corresponding PCR products are shown on the gel with the lanes labelled accordingly.

## Conclusions

The genome of *S. goldiniensis* ATCC 21386 was sequenced using a hybrid approach to yield a high-quality draft genome with ~99% of the genome on a single contig through a k-mer dependant assembly using SPAdes [9], followed by a scaffold-based final assembly with MeDuSa [13]. This allowed prediction of a biosynthetic gene supercluster to be identified from the main genome contig and the organisation of the supercluster to be confirmed by PCR. This genome sequence provides a springboard for further study of this strain and a basis for a formal taxonomic description of *S. goldiniensis* var. *goldiniensis* ATCC 21386.

## Materials and Methods

### Whole Genome Sequencing *Streptomyces goldiniensis*

Genomic DNA was extracted according to Kieser *et al*, [14] from cultures grown in GYM medium (DSMZ medium 65; www.dsmz.de). Nanopore sequencing was performed using the Nanopore™ 1D ligation protocol with MinION SPOT ON MK1 R9 flow cells. Raw data was converted using MinKnow base calling software. Illumina platform data was provided by Microbes NG (Birmingham, UK) from the HiSeq 2500 sequencing platform. Reads were trimmed using Trimmomatic 0.30 [15] with quality cut off of Q15. PacBio sequencing was provided by (Nu-omics, University of Northumbria, UK) using the Sequel instrument with contigs assembled in HGAP4.

The genome was assembled using SPAdes with combined data from all three platforms [9]. AutoMLST [16] was used to identify *S. bottropensis* ATCC 25435 (Taxonomy ID: 1054862) as the closest neighbour for scaffold-based assembly using MeDuSa [13] and quality analysis performed using QUAST [17]. Prokka was used to annotate the genome of *Streptomyces goldiniensis* [18] and is available on Genbank (Bioproject PRJNA602141). Identification of biosynthetic gene clusters was performed using the antiSMASH pipeline (bacterial version 5.0.0) [10]. The position of the putative aurodox BGC within the larger 273 Kbp ‘supercluster’was confirmed via PCR. The following oligonucleotide primer sequences were used;

clusterposcheckAF 5’-CCAGACGCAGGTCCGCTTCGGACG-3’;
clusterposcheckAR 5’-CCATCGTGGGGATCGCAG-3’;
clusterposcheckBF 5’-AGGATGTTCCAGTCGGCTCTCACTCCG-3’; clusterposcheckBR 5’-CGAGGTCGCCCGGCATGTGGA-3’.

PCR products were visualised using agarose gel-electrophoresis, and sequence specificity was confirmed by band excision followed by Sanger sequencing provided by Eurofins genomics (Luxembourg).

## Funding Information

We would like to thank the University of Strathclyde and University of Glasgow for jointly funding the PhD of REM. PAH would also like to acknowledge funding from BBSRC (BB/T001038/1 and BB/T004126/1) and the Royal Academy of Engineering Research Chair Scheme for long term personal research support (RCSRF2021\11\15). AJR & PAH would like to acknowledged funding from MRC (MR/V011499/1).

## Funding Statement

The funders had no role in study design, data collection and interpretation, or the decision to submit the work for publication

## Acknowledgements

We would also like to thank Professor Iain S. Hunter (University of Strathclyde) and Professor Matt Hutchings (John Innes Centre) for helpful discussions.

## Notes

https://www.ncbi.nlm.nih.gov/bioproject/PRJNA602141/

### Competing Interest Statement

The authors have declared no competing interest.

### Summary of Updates

Additional reference added describing the disection of the aurodox biosynthetic pathway. McHugh RE, Munnoch JT, Braes RE, Giard JM, Taladriz-Sender A, et al. Biosynthesis of aurodox, a Type III secretion system inhibitor from Streptomyces goldiniensis. BioRxiv. Epub ahead of print 2022. DOI: 10.1101/2022.02.14.480342. https://www.biorxiv.org/content/10.1101/2022.02.14.480342v1

